# Telomeric heterochromatin acts in *trans* to promote meiotic centromere assembly via Aurora B kinase recruitment

**DOI:** 10.64898/2026.06.06.730608

**Authors:** Haitong Hou, Ana Lopez Morales, Ying Liu, Julia Promisel Cooper

## Abstract

During meiosis, chromosomes face a paradox: the machinery that ensures reductional chromosome segregation also destabilizes centromeres by dismantling kinetochores, risking chromosome missegregation. Here we show how cells resolve this crisis through an unexpected activity of the telomere bouquet. We demonstrate that the bouquet transfers heterochromatin components to pericentromeres, which in turn recruit the Aurora B kinase to direct centromere reassembly. The heterochromatin protein Swi6^HP1^ relocates from telomeres to centromeres, enabling Haspin kinase-dependent phosphorylation of histone H3 and consequent enrichment of the chromosomal passenger complex, which includes the Aurora B kinase. Aurora B then phosphorylates core centromere proteins, including CenpA and CenpC, to promote kinetochore reassembly. Phosphomimetic mutants of CenpA or CenpC bypass the telomere-heterochromatin-Haspin pathway, demonstrating that Aurora B-mediated phosphorylation is sufficient for reassembly. This function is conserved in mitotically proliferating cells subjected to centromere dismantlement. Our findings establish a safeguarded system that couples meiotic nuclear architecture to centromere identity and reveal a fundamental role for the Aurora B kinase in centromere assembly, beyond its canonical function in correcting kinetochore-spindle attachment errors.

## Introduction

While centromeres and telomeres are structurally, functionally and spatially distinct through most cell cycle stages, their transient proximity in meiotic prophase promotes the reassembly of centromeres that have become dismantled through the action of meiosis specific factors [1, 2]. The mechanistic underpinnings of this ability of telomeres to promote centromere assembly have remained mysterious. Centromeres are chromosomal loci that serve as the assembly sites for kinetochores, the multi-protein complexes that allow spindle attachment and chromosome segregation in mitosis and meiosis. In most eukaryotes, including mammals and the fission yeast *Schizosaccharomyces pombe* (*S. pombe*), centromere positioning is epigenetically determined by the histone H3 variant CenpA. CenpA regions are typically flanked by pericentric heterochromatin (HC) characterized by histone H3 di- or tri-methylated at lysine 9 (H3K9me) bound by Heterochromatin Protein 1 proteins (HP1 in mammals; Swi6 in *S. pombe*). Thus, all three of the *S. pombe* centromeres are built upon unique central core sequences harboring CenpA flanked by repetitive pericentromeric HC. These features lend the ability to analyze the epigenetic features of centromeres in a genetically tractable model system.

Telomeres, comprising tandem G-rich repeats bound by shelterin proteins [3–5], generally prevent chromosome ends from being treated as damage-induced DNA double-strand breaks, averting chromosome end-fusion, checkpoint activation, and telomere entanglement [6, 7]. In meiotic prophase, telomeres cluster at the nuclear envelope (NE) to form a conserved structure known as the telomere bouquet [8]. The fission yeast bouquet localizes beneath the transmembrane LINC (Linker of Nucleoskeleton and Cytoskeleton) complex, which contacts the cytosolic spindle pole body (SPB), the yeast counterpart of the centrosome [9]. This is also the site where centromeres are anchored in mitotically proliferating cells [10]; importantly, a subset of telomeres and centromeres co-reside in this region during bouquet formation [2].

Centromeres remain remarkably stable over successive generations in proliferating cells [11, 12]. In meiosis, however, centromeres become precarious [2] as they are prone to dismantlement by Spo11 and Rec8, two key proteins that control foundational meiotic processes [1]. This dismantlement comprises loss of the entire kinetochore (including Cnp1, the *S. pombe* CenpA) as well as the pericentric HC, hereafter referred to as kinetochore loss, ‘KT loss’. KT loss is manifest specifically in the absence of the telomere bouquet or of HC, indicating that telomeric and HC factors protect centromeres from KT loss, likely through promoting reassembly [2]. Notably, the requirement for HC to reassemble dismantled meiotic kinetochores contrasts with the dispensability of HC for maintenance of CenpA chromatin through mitotic cell divisions; endogenous Cnp1 remains intact over many generations in the absence of the H3K9 methyltransferase Clr4^Suv39^ or Swi6^HP1^ [13].

Our observation that proximity between the bouquet and centromeres is important for meiotic kinetochore reassembly indicates that telomeres provide a specialized, assembly-promoting, microdomain [2]. Contiguous HC has been implicated in the establishment of Cnp1 chromatin during *de novo* centromere formation on newly introduced DNA [13] and during neocentromere formation following centromere excision [14]. However, the mechanism by which telomeres and HC promote reassembly of meiotic centromeres remains unknown. Here, we show that telomeric HC seeds pericentromeric HC in *trans*, leading to recruitment of the Aurora B kinase complex. Aurora B dependent phosphorylation of core kinetochore proteins then restores centromere integrity, revealing a mechanism by which nuclear architecture directs the re-establishment of centromere identity.

## Results

### Telomeric HC facilitates meiotic centromere assembly

At the onset of meiotic prophase, telomeres form the bouquet by moving to the LINC region of the NE prior to full centromere release from this region. This period of transient colocalization likely allows the clustered telomeres to establish a microenvironment that promotes centromere assembly [2]. To identify the relevant components of this microenvironment, we focused first on telomeric HC, as it harbors factors shared with pericentric HC. A point mutation in the shelterin protein Poz1 (Poz1-W209A) specifically disrupts telomeric HC in mitotically proliferating cells without compromising HC at other loci or canonical telomere functions, as indicated by normal telomere length [15]. We confirmed that bouquet formation remains robust in *poz1-W209A* cells (Figure 1A), allowing us to determine whether a bouquet lacking telomeric HC can promote centromere reassembly. We filmed live meiosis, using endogenously GFP-tagged Cnp3 (*S. pombe* CenpC) as a kinetochore marker along with marked chromatin (mCherry-tagged histone Pht1^H2AZ^), microtubules (mCherry-Atb2^α-Tubulin^), and SPBs (Sid4-mCherry) to indicate cell cycle stage [1]. Chromosomes with KT loss lack Cnp3-GFP and consequently remain unsegregated [1]. Quantitation revealed that *poz1W209A* confers KT loss at a level comparable to that imposed by disrupting the bouquet (*bqt1Δ*) or genome-wide HC (*clr4^Suv39^Δ* and *swi6^HP1^Δ*; Figure 1B, C). Notably, this defect is rescued by deletion of the genes encoding the meiosis-specific factors that dismantle centromeres, Spo11 or Rec8 (Figure 1D). Thus, telomere HC is required for the reassembly of kinetochores that have been compromised by the meiotic machinery, but is dispensable for maintenance of unperturbed meiotic kinetochores.

**Figure 1.**
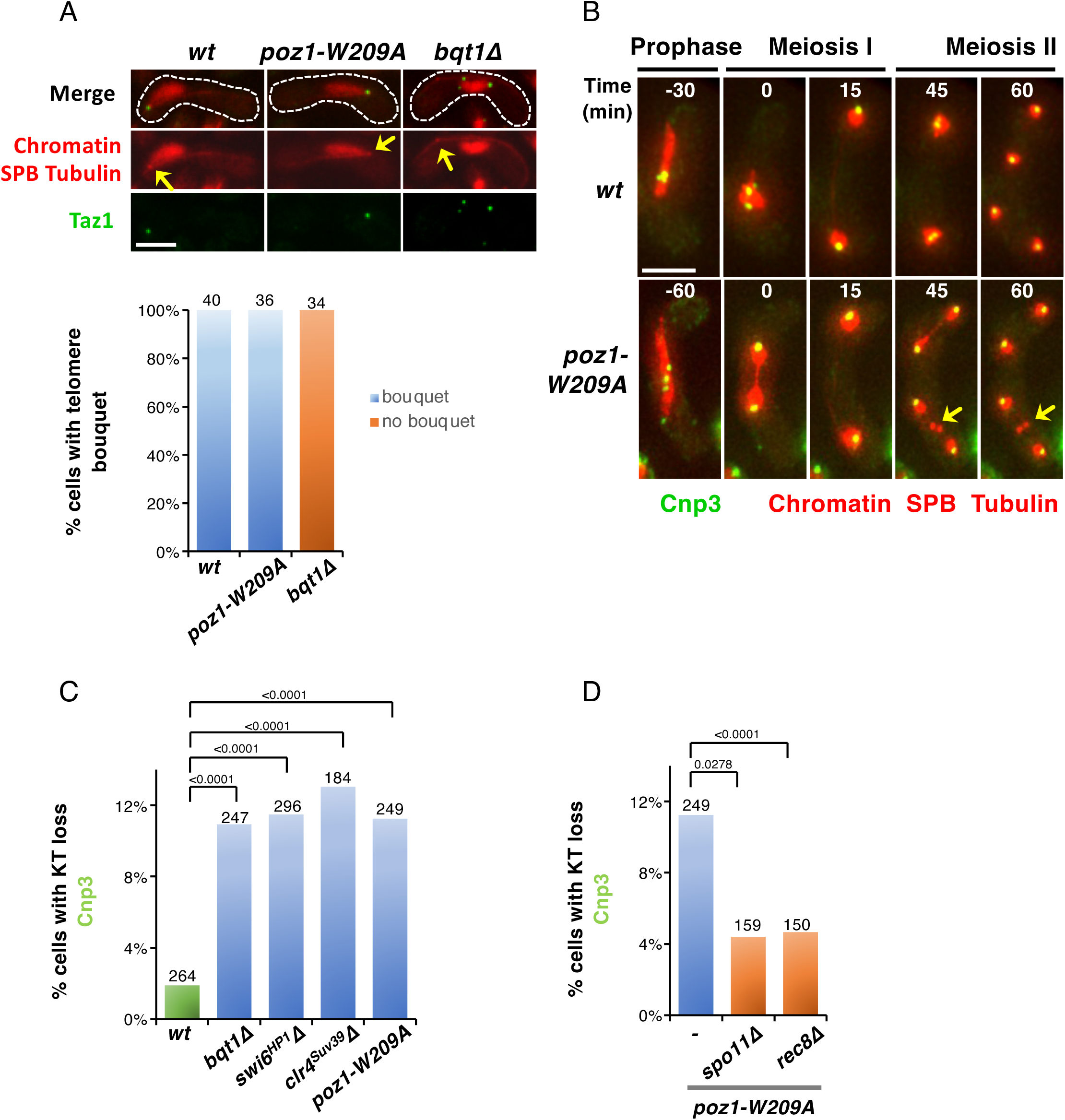
Telomeric HC promotes meiotic centromere assembly. **A.** Representative images (upper panel) and quantitation (lower panel) of bouquet formation in meiotic prophase. Telomeres (Taz1-GFP), SPBs (Sid4-mCherry), chromatin (Pht1^H2AZ^-mCherry) and microtubules (mCherry-Atb2^α-Tubulin^) are visualized. Arrow indicates the SPB. While *bqt1Δ* abolishes bouquet formation, *poz1-W209A* cells exhibit no defects in bouquet formation. Scale bar = 5 μm. The total number of meiotic prophase cells analyzed is indicated above each bar. **B.** Representative films of KT loss. KTs (Cnp3-GFP), chromatin (Pht1^H2AZ^-mCherry), SPBs (Sid4-mCherry) and microtubules (mCherry-Atb2^α-Tubulin^) are visualized. t = 0 corresponds to the first frame in meiosis I. Arrows indicate unsegregated chromosomes lacking KTs. Scale bar = 5 μm. **C-D**. Quantitation of KT loss from films as described for **B**. *poz1-W209A* cells show KT loss rates similar to *bqt1Δ*, *swi6^HP1^Δ*, and *clr4^suv39^Δ* cells; this is rescued by deletion of *spo11+* or *rec8+*. n values are indicated above each bar. P values, determined by two-tailed Fisher’s exact tests, are indicated above the brackets.

### The telomere bouquet promotes assembly of pericentromeric HC

How does telomeric HC promote the reassembly of centromeres? One possibility is that HC components such as Swi6^HP1^ migrate from telomeres to centromeres when they colocalize during bouquet formation, thereby restoring pericentromeric HC, which would in turn promote the reassembly of Cnp1 nucleosomes. An alternative model would invoke direct restoration of CenpA assembly by telomeric HC.

To explore these possibilties directly, we assessed the transfer of telomeric HC to pericentromeres using a zygotic meiosis assay (Figure 2, Supplementary Figure S1), in which meiosis is triggered immediately after mating two haploid strains of opposite mating types. During meiotic prophase in a wild type (*wt*) setting, we find that the endogenously tagged kinetochore marker Mis6^CenpI^-GFP, inherited from only one mating partner, colocalizes with Swi6^HP1^, inherited only from the opposite partner (Supplementary Figure S1A, B). As expected, this pericentric HC assembly depends on the RNAi pathway [16], as none of the Mis6^CenpI^ foci colocalize with Swi6^HP1^ when both parental strains lack Dcr1 (Figure 2). To determine whether HC can transfer from telomeres to pericentromeres, we engineered a cross between (1) a strain that lacks HC entirely (*swi6^HP1^τ<*) and harbors endogenously GFP-tagged Mis6^CenpI^, and (2) a *dcr1τ<* strain that harbors telomeric HC but lacks pericentric HC and carries endogenously GFP-tagged Swi6^HP1^ (Figure 2A). The zygotes derived from this cross show Swi6^HP1^ localization to most Mis6^CenpI^ foci during meiotic prophase (Figure 2B, C). Therefore, pericentric HC has assembled during meiotic prophase, despite the absence of pericentric HC in either parental strain. Remarkably, this meiotic assembly of pericentric HC depends on the bouquet; when both mating partners lack Bqt1, the number of Mis6^CenpI^ foci lacking Swi6^HP1^ increases substantially (Figure 2B, C). Thus, the bouquet promotes passage of Swi6^HP1^ from telomeres to pericentromeres, demonstrating a powerful *trans* effect of telomeric HC on pericentric HC assembly.

**Figure 2.**
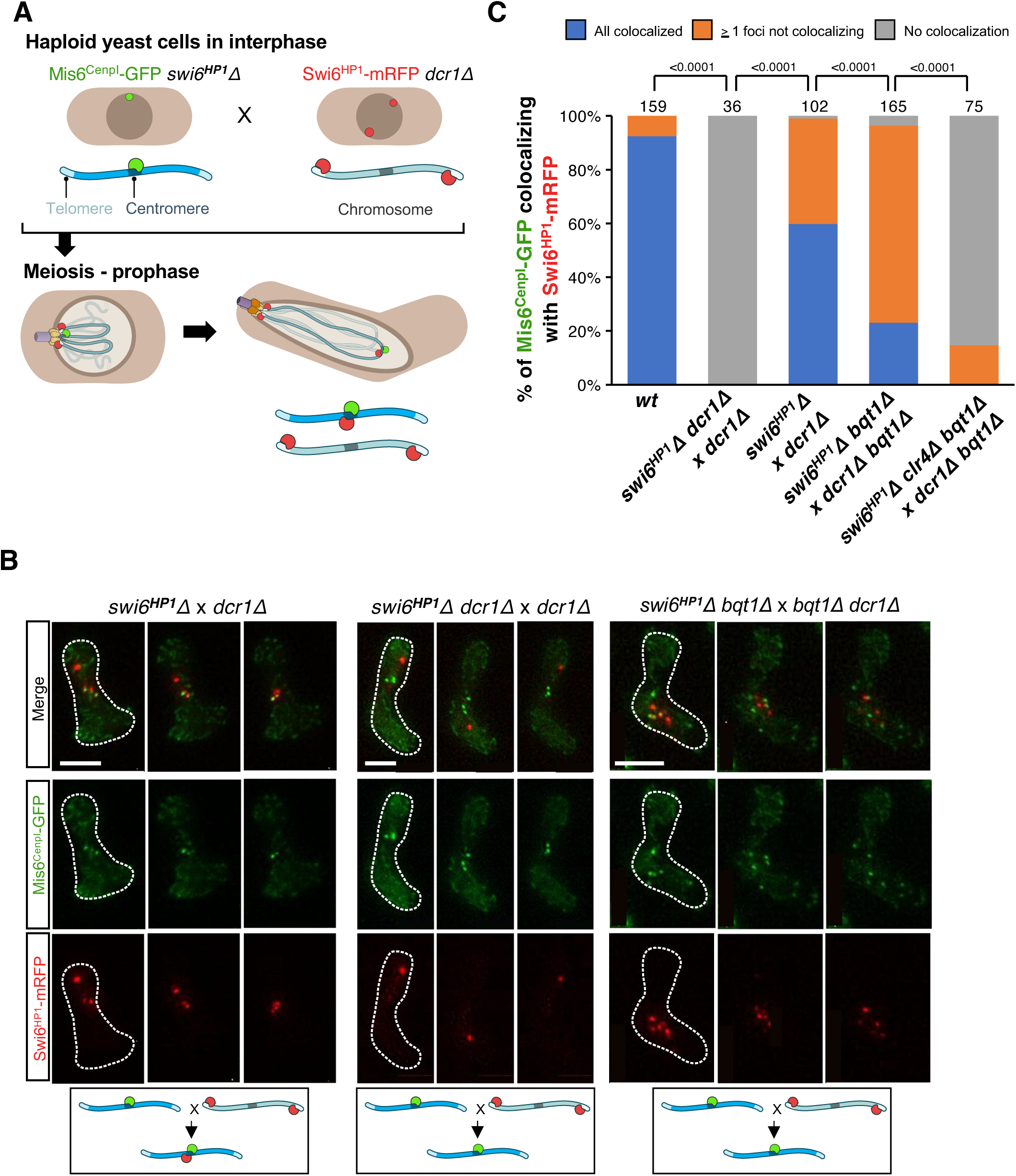
The telomere bouquet promotes reassembly of pericentromeric HC. **A.** Schematic of experimental setup for monitoring meiotic pericentromeric HC assembly. Haploid cells of opposite mating types, one expressing Mis6^CenpI^-GFP and the other expressing Swi6^HP1^-mRFP, are mated. The diagram exemplifies the *wt* scenario. Colocalization of the two markers during meiotic prophase is quantified. **B-E**. Frames from representative films of meiotic prophase with tagged proteins as described in **A**. Localization of Swi6^HP1^-mRFP and Mis6^CenpI^-GFP is monitored across the indicated genetic backgrounds. Scale bar = 5 μm. **F.** Quantitation of colocalization of Swi6^HP1^-mRFP and Mis6^CenpI^-GFP. Colocalization is reduced in *the absence of Dcr1 or of the bouquet*. n values are indicated above each bar. P values, determined by two-tailed Fisher’s exact tests, are indicated above the brackets.

In an independent zygotic HC transfer assay, one mating partner harbored Swi6^HP1^ endogenously tagged with slow-folding red fluorescent protein (sfRFP) [17, 18], while the other partner habored untagged Swi6^HP1^ (Supplementary Figure S1C, D). In both partners, mCherry-Atb2^α-Tubulin^ was expressed to visualize microtubules, while a fluorescent repressor array (*lacO-lacI-GFP*) inserted within the innermost repeats of pericentromere I (*Imr1*) [19] allowed us to monitor Swi6^HP1^ flanking the centromere. A characteristic cytoplasmic microtubule pattern [20, 21] and the appearance of two unpaired *Imr1*-GFP foci on homologous chromosomes indicate a meiocyte in early meiotic prophase.

In an otherwise *wt* setting, Swi6^HP1^-sfRFP foci localize adjacent to both *Imr1*-GFP foci (Supplementary Figure S1D), reflecting intact pericentric HC. Since sfRFP requires approximately 11 hours to mature, the Swi6^HP1^-sfRFP signal at the *Imr1* locus inherited from the partner harboring untagged Swi6^HP1^ must originate from pre-existing Swi6^HP1^-sfRFP provided by its mating partner. This indicates that pericentromeric Swi6^HP1^ molecules can exchange dynamically, allowing Swi6^HP1^ from one mating partner to assemble at the pericentrome of the other partner at an early stage of meiosis. When both mating partners lack Dcr1, the zygotes fail to exhibit Swi6^HP1^-sfRFP signals on either *Imr1*-GFP focus (Supplementary Figure 1D), consistent with the established role of the RNAi pathway in pericentromere HC assembly [2, 16].

Crucially, in over 80% of *bqt1Δ* or *poz1W209A* cells, Swi6^HP1^-sfRFP signals are absent from one of the two *Imr1*-GFP dots (Supplementary Figure S1D), indicating that a bouquet harboring telomeric HC is required for Swi6^HP1^ assembly at meiotic centromeres; a defective bouquet lacking telomeric HC fails to confer Swi6^HP1^ assembly. Collectively, these observations demonstrate that Swi6^HP1^ molecules move from the telomere bouquet to pericentromeres upon bouquet formation. We surmise that this HC transfer mediates the ability of telomeres to promote pericentromeric HC assembly, which subsequently stimulates the reassembly of CenpA chromatin.

### HC recruits Aurora B kinase to facilitate meiotic centromere assembly

Given the key role of HC in telomere bouquet-mediated centromere assembly, we sought to identify downstream factors important for this process. Pericentric HC recruits chromosome segregation factors including cohesins and Shugoshin [15–17], and in mitotically proliferating cells recruits Haspin kinase (Hrk1) through interactions with Swi6^HP1^ and the cohesin-associated protein Pds5 [22, 23]. Haspin phosphorylates histone H3 at threonine 3 (H3T3ph) [22, 24, 25], which in turn recruits the chromosomal passenger complex (CPC) (Figure 3A). The CPC catalytic subunit, the Aurora B kinase (Ark1), promotes chromosome biorientation by correcting erroneous KT-spindle attachments.

**Figure 3.**
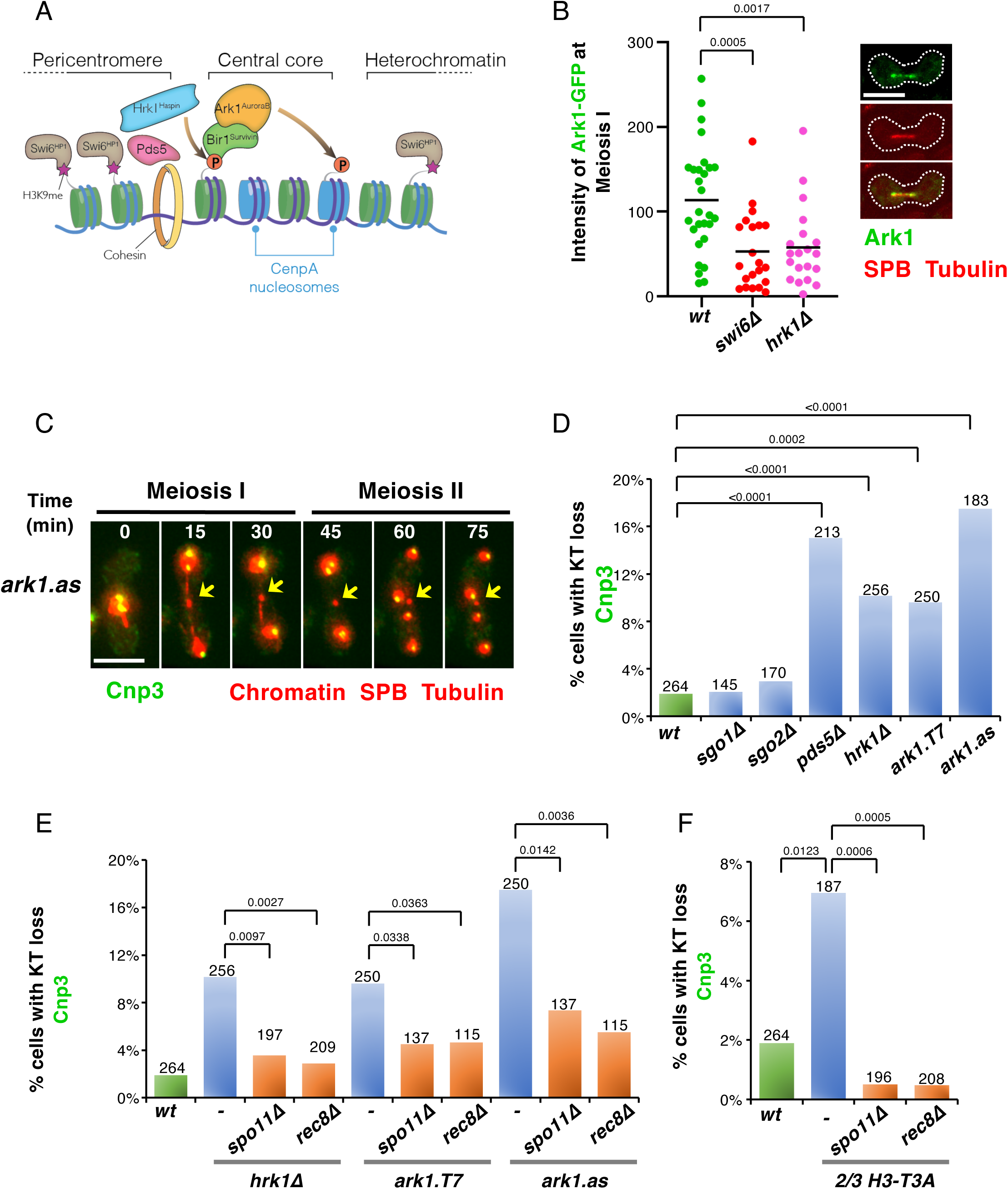
HC promotes kinetochore reassembly by recruiting Aurora B. **A.** Diagram illustrating HC-mediated recruitment of Ark1^AuroraB^ to centromeric chromatin. The core centromere, marked by Cnp1^CenpA^ nucleosomes, is flanked by pericentric HC harboring H3K9me and Swi6^HP1^. This HC recruits Ark1^AuroraB^ via the cohesin associated protein Pds5 and Hrk1-dependent H3T3 phosphorylation, both of which are promoted by interactions with Swi6^HP1^. **B.** Quantitation of Ark1localization along spindles during metaphase I. Representative images of cells at this stage are also shown. Ark1-GFP, SPBs (Sid4–mCherry), and microtubules (mCherry–Atb2^α-Tubulin^) are visualized. Both *swi6^HP1^Δ* and *hrk1Δ* cells show decreased centromeric localization of Ark1-GFP. Scale bar = 5 μm. P values, determined by two-tailed Fisher’s exact tests, are indicated above the brackets. **C.** Frames from representative film of KT loss in *ark1.as* cells. KTs (Cnp3–GFP), chromatin (Pht1^H2AZ^–mCherry), SPBs (Sid4–mCherry), and microtubules (mCherry–Atb2^α-Tubulin^) are visualized. t = 0 is the first frame in meiosis I. The arrows indicate unsegregated chromosomes that lack KTs. Scale bar = 5 μm. **D-F**. Quantitation of KT loss as in **C**. All mutations that compromise recruitment of the CPC to pericentromeric HC cause KT loss **(D)** in a manner that depends on the presence of Spo11 and Rec8 **(E)**. Mutating histone H3-T3 to Alanine at two of three histone H3 alleles causes KT loss, which is rescued by *spo11+* or *rec8+* deletion **(F)**. The total number of meiocytes analyzed is indicated above each bar. P values, determined by two-tailed Fisher’s exact tests, are indicated above the brackets.

The existence and roles of the HC-Haspin-CPC pathway in meiosis have been less clear. We find that Ark1-GFP appears at centromeres from prometaphase to metaphase I (Supplementary Figure S2A) [26]. In meiocytes lacking either Swi6^HP1^ or Hrk1, centromeric Ark1-GFP signal is significantly reduced (Figure 3B). Residual Ark1-GFP signal likely reflects a second recruitment pathway mediated by the Shugoshin Sgo2 [27]. In mitosis, CPC recruitment can also occur through a Bub1-Histone H2A-S121ph-Sgo2 pathway upon activation of the spindle assembly checkpoint (SAC) [28]. Consistent with a similar contribution in meiosis, deletion of *sgo2^+^* reduces Ark1-GFP at centromeres, and combined loss of Hrk1 and Sgo2 nearly abolishes the signal (Supplementary Figure S2B).

To test whether CPC recruitment mediates the role of HC in facilitating meiotic centromere assembly, we examined the KT loss phenotype in an array of mutants along this pathway. *pds5Δ* and *hrk1Δ* cells show KT loss at levels comparable to those seen in the absence of the bouquet or HC (Figure 3C, D). Epistasis analysis revealed that simultaneously deleting *hrk1^+^* and either *bqt1^+^* or *swi6^HP1+^* results in levels of kinetochore loss similar to the single mutants (Supplementary Figure S2C). Thus, the bouquet, HC and Haspin act through a common pathway.

Crucially, as is the case in *bqt1Δ* cells, KT loss in *hrk1Δ* cells is meiosis-specific, and rescued by deletion of *spo11^+^*or *rec8^+^* (Figure 3E). Thus, Haspin is specifically required to reassemble centromeres that have been dismantled by Spo11 and Rec8.

Consistent with a role for the HC-Haspin-H3T3ph-CPC pathway in kinetochore reassembly, mutation of H3T3 to a non-phosphorylatable alanine at two of the three histone H3 loci (*hht1-T3A hht3-T3A* or ‘*2/3 h3-T3A*’) causes KT loss (Figure 3F). Indeed, the KT loss rate in *2/3 h3-T3A* cells (7%) is comparable to that seen in *hrk1 Δ* cells (10.1%), consistent with H3-T3 being the primary Haspin substrate. As in *hrk1Δ* cells, this phenotype is rescued by deleting *spo11^+^* or *rec8^+^* (Figure 3F). These results indicate that the HC-mediated enrichment of Aurora B activity promotes meiotic centromere reassembly but is dispensable for maintenance of unchallenged centromeres.

To directly test the role of the Aurora B kinase in centromere reassembly, we examined two hypomorphic alleles: *ark1-T7*, a temperature-sensitive (*ts*) allele [27], and *ark1-as*, an ATP-analog-sensitive (*as*) allele. The *ark1-T7* mutant exhibits meiotic KT loss at the semi-permissive temperature of 30°C. Consistent with reduced kinase activity, *ark1-as* cells show KT loss even without the addition of ATP-analog inhibitor (Figure 3C, D). KT loss in both *ark1-T7* and *ark1-as* cells is substantially rescued by deletion of *spo11^+^* or *rec8^+^* (Figure 3E), indicating that Aurora B activity is specifically required to restore meiotically dismantled kinetochores.

Deletion of *sgo2^+^* does not cause meiotic KT loss (Figure 3D), suggeswwng that the pool of HC-enriched Aurora B kinase is specifically required for centromere reassembly. This is consistent with the notion that centromeric Sgo2 localization depends on intact centromeres and is therefore lost upon dismantlement, whereas HC assembly can occur independently of centromere identity.

During meiosis, pericentromeric Swi6^HP1^ recruits not only Hrk1 but also the meiosis-specific Shugoshin Sgo1, which protects cohesion of sister chromatids at Meiosis I [29, 30]. We examined *sgo1Δ* cells and observed no increase in KT loss rate above the wt level (Figure 3D). Thus, Sgo1 is not required for kinetochore reassembly.

### Ectopically targeting Haspin or CPC to centromeres bypasses the requirement for HC

If HC facilitates meiotic centromere assembly primarily by recruiting the CPC, can the requirement for HC be bypassed by artificially tethering the CPC to centromeres? Fusions between two tandem H3K9Me-binding chromodomains (derived from Swi6^HP1^) and either the CPC component Bir1^Survivin^ (Bir1^Survivin^-CD) [27] or Hrk1 (Hrk1-CD) [31] have been used successfully to force CPC localization to centromeres. In *swi6^HP1^Δ* cells, residual pericentromeric H3K9me is sufficient to interact with the chromodomains (CD) and recruit these fusion proteins (Figure 4A, B), restoring the function of the CPC in correcting erroneous kinetochore attachment [31]. Using this approach, we find that ectopic recruitment of Bir1^Survivin^-CD partially rescues the meiotic KT loss phenotype of *swi6^HP1^Δ* cells (Figure 4C). In contrast, Bir1^Survivin^-CD failed to rescue the phenotype in *clr4^suv39^Δ* cells, in which H3K9me is abolished, leaving no mark for CD recruitment. Similarly, a fusion between the CDs and Hrk1, but not a kinase-dead (KD) allele [31], rescues the KT loss phenotype of *swi6^HP1^Δ* cells. As expected, Hrk1-CD was also unable to rescue the phenotype in *clr4^suv39^Δ* cells (Figure 4C). Therefore, recruitment of the CPC mediates the role of HC in promoting meiotic centromere reassembly.

**Figure 4.**
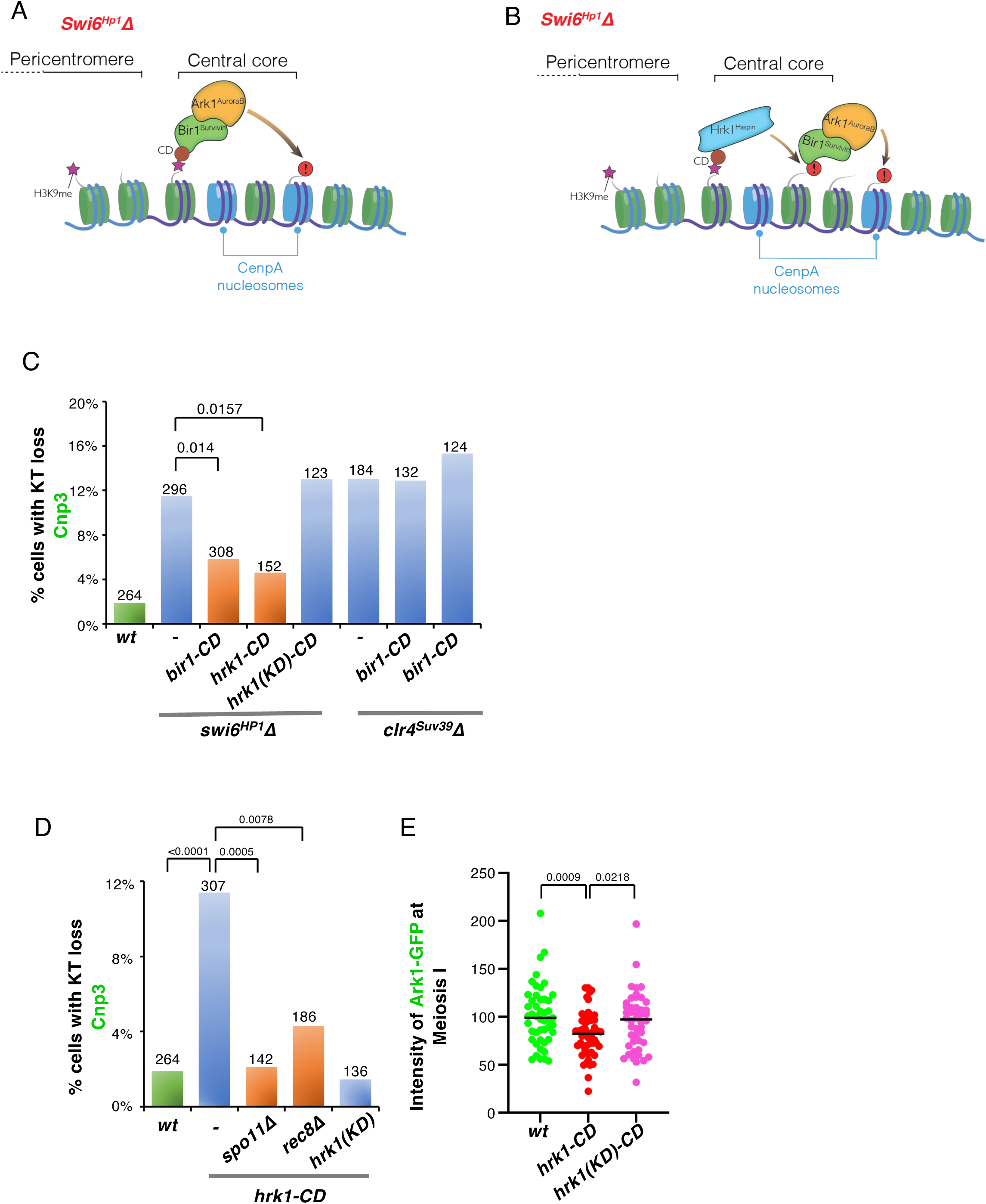
Ectopically targeting CPC to centromeres bypasses the requirement for HC for kinetochore reassembly. **A-B.** Schematic of CPC targeting system in *swi6^HP1^Δ* cells. Bir1^Survivin^ is fused with H3K9me-binding chromodomains (CDs) **(A),** or CDs are fused with Hrk1^Haspin^, which locally phosphorylates H3T3 to recruit the CPC via Bir1^Survivin^ binding **(B)**. **C-D.** Quantitation of KT loss. Bir1-CD and Hrk1-CD fusions rescue KT loss in *swi6^HP1^Δ* cells but not in *clr4 ^suv39^Δ* cells. The catalytically dead Hrk1(KR)-CD fails to rescue KT loss in *swi6^HP1^Δ* cells **(C)**. In a *wt* background, Hrk1-CD, but not Hrk1(KR)-CD, induces KT loss in a manner that depends on the presence of Spo11 and Rec8 **(D)**. The total number of meiocytes analyzed is indicated above each bar. P values, determined by two-tailed Fisher’s exact tests, are indicated above the brackets. **E.** Quantitation of Ark1-GFP intensity at metaphase I. In a *wt* background, Hrk-CD, but not Hrk1(KR)-CD, confers reduced Ark1-GFP levels at centromeres. P values, determined by two-tailed Fisher’s exact tests, are indicated above the brackets.

Curiously, ectopic expression of Hrk1-CD in a *wt* background induced KT loss (Figure 4D). This phenotype was rescued by deletion of *spo11^+^* or *rec8^+^* (Figure 4D), indicating that it stems from an inability to reassemble kinetochores dismantled by these factors. We attribute this phenotype to titration of the CPC away from the centromere region and towards other HC regions, presumably including the mating type locus and subtelomeres. While all HC is lost in *swi6^HP1^Δ* cells, the intact HC in *wt* cells may nonselectively confer recruitment of Hrk1-CD to these non-centromeric HC regions, titrating Ark1 away from centromeres. Indeed, we observed reduced Ark1-GFP intensity at centromeres in meiosis I upon expression of Hrk1-CD (Figure 4E). As a control, ectopic expression of the kinase-dead Hrk1 (KD)-CD, which cannot install additional H3T3ph sites, does not affect centromeric Ark1-GFP signal intensity (Figure 4E), consistent with its inability to dismantle centromeres (Figure 4D).

Collectively, our data demonstrate that HC promotes meiotic centromere assembly by recruiting Haspin, which enriches the CPC at centromeres through H3T3 phosphorylation. The centromere-localized activity of Aurora B kinase is in turn crucial to counteract KT dismantlement by Spo11 and Rec8.

### Compromised CPC exacerbates Spo11-mediated centromere dismantlement in proliferating cells

Previously, we found that in mitotically proliferating cultures, ectopic expression of Spo11 (under control of the constitutive *adh1* promoter) causes complete KT loss in approximately 5% of cells and reduces the overall amount of Cnp1 at all centromeres, including those on properly segregated chromosomes [1]. We therefore asked whether centromere reassembly in proliferating cells is promoted by centromeric CPC. Mitotically proliferating cells lacking Swi6^HP1^ or Hrk1 exhibit few lagging chromosomes devoid of the kinetochore marker Ndc80-2xGFP (Supplementary Figure S3A, B). Thus, the CPC-mediated centromere assembly pathway is dispensable when centromeres are not being actively dismantled by meiotic proteins. In contrast, the frequency of lagging chromosomes lacking kinetochore signals increases significantly when *swi6^HP1^Δ* or *hrk1Δ* cells are challenged by Spo11 expression (Supplementary Figure S3A, B). Thus, the HC-Haspin-CPC pathway possesses the capacity to reassemble dismantled centromeres not only in meiosis, but also in mitotically proliferating cells.

### Aurora B kinase promotes centromere assembly by phosphorylating CenpA and CenpC

How does the CPC promote centromere assembly? To address this, we sought to identify crucial substrates of the Aurora B kinase. Given that all centromeric proteins are completely erased from dismantled centromeres in *ark1* mutants, phosphorylation of core structural components of the kinetochore, such as CenpA and CenpC, emerged as potential candidates. Indeed, Aurora B kinase phosphorylation motifs are conserved across eukaryotes within the N-terminal domain of CenpA, despite the low overall sequence conservation of this region (Figure 5A) [32].

**Figure 5.**
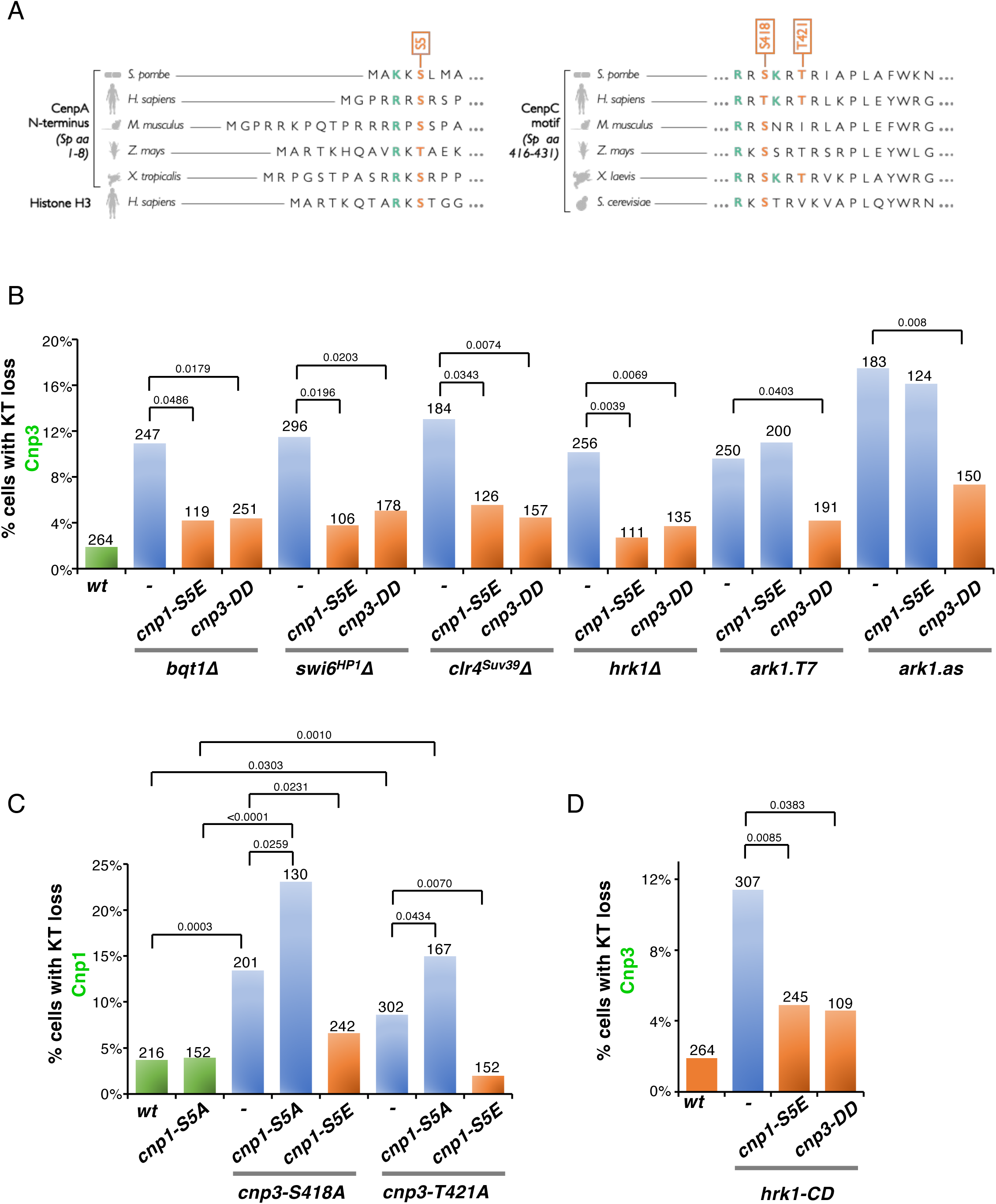
Phosphorylation of CenpA and CenpC by Aurora B kinase promotes meiotic centromere assembly. **A.** Sequence alignment showing conserved Aurora B kinase phosphorylation sites at the N-terminus of CenpA and within the CenpC motif. Phosphorylated residues within consensus motifs are shown in red; additional key consensus motif residues are shown in green. **B-D.** Quantitation of meiotic KT loss as exemplified in Figure 1B and 3C. Phosphomutations of Cnp3 (S418A or T421A) elevate meiotic KT loss, which is further increased when combined with the Cnp1-S5A mutation **(B)**. The phosphomimetic mutations Cnp1-S5E) and Cnp3-DD (S418D T421D) rescue KT loss in *bqt1Δ*, *swi6^HP1^Δ*, *clr4^suv39^Δ, hrk1Δ*, *ark1.T7*, and *ark1.as* mutants **(C)**, as well as in cells expressing *bir1-CD* and *hrk1-CD* **(D)**. The total number of meiocytes analyzed is indicated above each bar. P values, determined by two-tailed Fisher’s exact tests, are indicated above the brackets.

Mutation of the conserved Aurora B kinase site at serine 5 of endogenous Cnp1 to a non-phosphorylatable alanine (Cnp1-S5A) yields no growth defect nor enhanced sensitivity to the microtubule-destabilizing drug thiabendazole (TBZ) (Supplementary Figure S4A); likewise, chromatin immunoprecipitation (ChIP) shows similar levels of centromeric enrichment for Cnp1-S5A and *wt* Cnp1 (Supplementary Figure S4B). Remarkably, however, the phospho-mimetic mutant Cnp1-S5E, in which serine 5 is endogenously replaced by glutamate, is sufficient to rescue the centromere dismantlement phenotypes of *bqt1Δ*, *swi6^HP1^Δ*, *clr4^Suv39^Δ* and *hrk1Δ* cells (Figure 5B). Thus, Aurora B kinase-mediated phosphorylation of Cnp1 can promote meiotic centromere assembly. Nonetheless, meiotic *cnp1-S5A* cells do not exhibit increased centromere dismantlement compared to *wt* cells (Figure 5C). Hence, phosphorylation at sites additional to the Cnp1 N-terminus likely act redundantly to promote meiotic centromere assembly.

The essential centromere protein CenpC promotes centromere assembly by recruiting the Mis18 kinetochore assembly complex [33, 34]. The centromeric localization of CenpC depends on its conserved CenpC motif, which directly interacts with CenpA [35, 36] and is regulated by CDK1-mediated phosphorylation [37, 38]. Moreover, Aurora B kinase consensus motifs within the CenpC motifs are conserved across eukaryotes (Figure 5A); the function of modification at these sites remains to be determined. The putative Aurora B phosphorylation sites in *S. pombe* CenpC (Cnp3) are serine 418 and threonine 421. Like Cnp1-S5E, a phospho-mimetic allele of endogenous Cnp3 (S418D T421D, hereafter referred to as Cnp3-DD) rescues the KT loss phenotype of *bqt1Δ*, *swi6^HP1^Δ*, *clr4^suv39^Δ* and *hrk1Δ, ark1-T7* and *ark1-as* cells (Figure 5B). Thus, constitutive phosphorylation of either Cnp1 or Cnp3 is sufficient to rescue the meiotic KT loss phenotype induced by Spo11 and Rec8.

To address the necessity of Cnp3 phosphorylation for kinetochore reassembly, phosphomutant alleles were constructed. The phospho-deficient mutant Cnp3-S418A diminishes Cnp3-GFP intensity at centromeres in mitotically proliferating cells (Supplementary Figure S4B) and causes growth defects (Supplementary Figure S4A). This reduction in Cnp3 intensity is clearly distinct from KT loss, in which all centromere markers including Cnp1 are absent, but makes Cnp3-GFP unsuitable as a marker for quantifying KT loss. Nonetheless, mitotic Cnp1 levels show little change, as measured by ChIP (Supplementary Figure S4A), in either Cnp3-S418A and Cnp3-T421A backgrounds, allowing use of Cnp1-GFP as a KT loss marker. Both Cnp3 mutants cause meiotic KT loss (Figure 5C) and this is exacerbated when combined with *cnp1-S5A*, indicative of functional redundancy between phosphorylated Cnp1 and phosphorylated Cnp3.

Cnp3-T421A-GFP is also bright enough in both mitotically proliferating and meiotic cells to serve as a reliable marker. Cnp3 loss is induced by the *cnp3-T421A* mutant; this is further enhanced when combined with *cnp1-S5A* (Supplementary Figure S4D). Moreover, the KT loss phenotypes induced by *cnp3-S418A* and *cnp3-T421A* phospho-mutants, alone or in combination with *cnp1-S5A*, are rescued by deletion of *spo11+* or *rec8+*, confirming that these centromere defects are meiosis specific, and that these phosphosites are dispensable in the absence of active kinetochore dismantlement by Spo11 or Rec8 (Supplementary Figure S4E, F).

Thus, while phosphorylation of the CenpC motif of Cnp3 is important for meiotic centromere reassembly, phosphorylated Cnp1 acts redundantly to promote reassembly. A phospho-mimetic mutation in either protein is sufficient to bypass the requirement for the telomere bouquet, HC, and Haspin. Supporting this notion, Cnp1-S5E rescued the centromere dismantlement caused by the phospho-deficient mutants Cnp3-S418A or -T421A (Figure 5B, Supplementary Figure S4D), confirming the sufficiency of constitutive Cnp1 phosphorylation. In contrast to the suppression by Cnp1-S5E of KT loss in *bqt1τ<* or HC-deficient settings, Cnp1-S5E fails to suppress KT loss in cells harboring *ark1-T7* or *ark1-as* (Figure 5B). This implicates a pool of Aurora B kinase distinct from the fraction enriched by HC that contributes to centromere assembly. Indeed, the severity of centromere dismantlement in the *ark1-as* background is slightly greater than in mutants compromising the bouquet-HC-Haspin-H3T3ph pathway. As Cnp3-DD suppresses the hypomorphic Ark1 alleles (Figure 5B), the additional Aurora B substrates may promote kinetochore reassembly by stabilizing Cnp3 at centromeres.

## Discussion

Despite the remarkable stability of CenpA chromatin, the telomere bouquet is required to counteract centromere disassembly by key meiotic proteins [1, 2]. Here, we demonstrate that the heterochromatic nature of the telomere bouquet is crucial for this function, as a bouquet devoid of telomeric HC fails to promote centromere reestablishment. This process resembles *de novo* centromere formation and likely shares requirements with neocentromere formation.

Swi6^HP1^ relocates from telomeres to pericentric regions during meiosis, as evinced by our zygotic meiosis assays that show transfer of Swi6^HP1^ from the telomeres of one mating partner to the pericentromeres of another. This transfer depends on bouquet formation, indicating that the mere presence of Swi6^HP1^ in the nucleus is insufficient to ensure *de novo* pericentric HC assembly. Rather, the spatial concentration of HC components imparted by clustered telomeres creates a microenvironment that promotes efficient pericentromere assembly.

This trans-acting redistribution of HC components is notable in light of observations in *Drosophila melanogaster* [39], in which distinct HC domains, the pericentric regions and the rDNA, are spatially associated but remain compositionally distinct, connected by bridging interactions rather than by mixing. Together with that work, our findings raise the possibility that the telomere bouquet creates a specialized nuclear microdomain that permits regulated transfer or exchange of HC components under defined conditions, rather than wholesale mixing of HC states.

The mechanisms by which HC promotes nearby CenpA nucleosome incorporation have remained largely mysterious. Here we identify HC-mediated recruitment of the Aurora B kinase complex, via the Haspin-H3T3ph pathway, as a critical driver of meiotic centromere reassembly. While the canonical role of centromeric Aurora B is to eliminate erroneous kinetochore-microtubule attachments [40], our observations reveal an additional function in catastrophic scenarios of meiotic KT loss requiring *de novo* centromere assembly. Thus, whereas Polo kinase and CDK1 license cell cycle-coupled replenishment of CenpA following dilution due to DNA replication [33], Aurora B appears to function specifically upon total CenpA loss, when centromeric identity is abolished.

Aurora B is also required for centromere assembly in mitotically proliferating cells whose kinetochores are challenged by misexpression of Spo11 (Supplementary Figure S3). Therefore, the Aurora B pathway may also promote *de novo* centromere assembly on newly introduced DNA, as well as neocentromere formation at subtelomeric regions following the excision of an endogenous centromere [13, 14]. A recent model proposes that HC facilitates *de novo* centromere establishment on plasmids by tethering them to the endogenous centromeric microdomain beneath the LINC complex, where Cnp1 and possibly other assembly factors are enriched [34]. Aurora B is likely a critical component of this centromere assembly-promoting microdomain.

We propose a two-step model for meiotic centromere reassembly (Figure 6). First, during bouquet formation beneath the SPB, HC components including Swi6^HP1^ are transferred from telomeres to centromeres, restoring or reinforcing pericentric HC following disruption by Spo11 and Rec8. Second, Aurora B kinase is recruited to this HC domain, where its activity promotes reassembly of CenpA chromatin and kinetochore structure prior to meiosis I. In a bouquet-deficient setting, the last centromere to dissociate from the LINC region is left in a domain with neither pericentric HC from other chromosomes nor telomeres. Thus, these late-departing centromeres have no neighboring HC to impart Swi6^HP1^ transfer and no recourse following Spo11/Rec8-mediated dismantlement.

**Figure 6.**
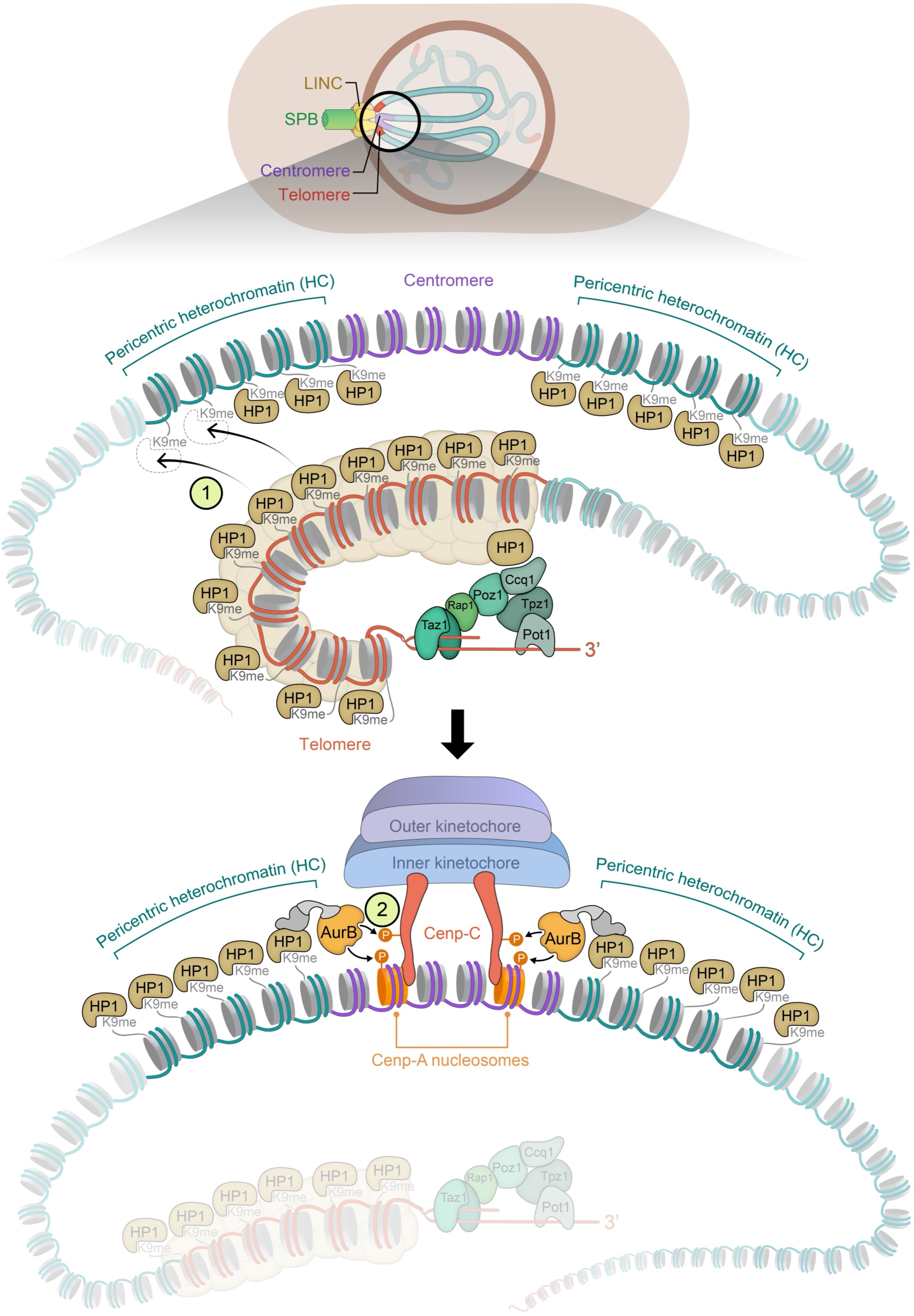
A two-step model for telomeric heterochromatin-mediated centromere assembly in meiosis. Step 1. During bouquet formation, telomeres transiently colocalize with centromeres, enabling the transfer of telomeric HC components such as Swi6^HP1^ to pericentromeric regions. This facilitates the reestablishment of pericentromeric HC that has been dismantled by the meiotic factors Spo11 and Rec8 [1]. Step 2. At the onset of meiosis I, Aurora B kinase is recruited to pericentric HC, where it phosphorylates core centromere proteins (CenpA and CenpC), promoting kinetochore reassembly. In the absence of the bouquet or HC, this pathway fails, leading to persistent KT loss.

We find that both the HC-Haspin-H3T3 and Bub1-H2AS121ph-Shugoshin pathways contribute to the localization of the CPC to meiotic centromeres (Supplementary Figure S2B). However, only the HC-Haspin pathway contributes to KT reassembly. We suggest that this reflects the distinct requirements of fully dismantled centromeres, which lack the kinetochore-dependent machinery needed to recruit Bub1 and Sgo2. In contrast, HC can be re-established independently of centromere identity, providing a platform for CPC recruitment via the HC-Haspin pathway.

Intriguingly, Sgo2 is also present at the ‘knob’, a region interior to subtelomeres, and its localization there depends on Bub1 [41]. However, our results suggest that this pool of Sgo2 is either insufficiently enriched or positioned too far from the telomere to support centromere reassembly.

How could the phosphorylation of CenpA and CenpC promote centromere assembly? Both direct and indirect phospho-mediated stabilization of these factors at centromeres may be involved. The function of the CenpA N-terminus harboring the putative Aurora B kinase phosphosites remains enigmatic [42–44]. Intriguingly, N-terminal truncation of *S. pombe* Cnp1 increases its centromeric level, implying that this region auto-inhibits its own deposition [45]. This finding invites a speculative analogy with a known regulatory mechanism: Aurora B kinase-mediated phosphorylation of the KT protein Dsn1 relieves its auto-inhibition in budding yeast and human cells, permitting kinetochore assembly [46–48]. By extension, Aurora B kinase-mediated phosphorylation of the CenpA N-terminal tail may liberate the histone-folding domain to productively engage with centromere assembly factors.

Aurora B kinase-mediated phosphorylation within the CenpC motif may strengthen both CenpC’s centromeric localization and its interaction with pre-existing CenpA. Indeed, the centromere localization of CenpC is markedly diminished when the phosphorylation site Cnp3-S418 is mutated (Supplementary Figure S4C). Consistently, mutating other residues (R416 and R420) within the Aurora B kinase consensus motif (Figure 5A) also reduces CenpC localization at centromeres [37, 49]. CenpA deposition is mediated by its chaperone, HJURP (Scm3 in *S. pombe*), which is recruited to the centromeric region in a CenpC-dependent manner during G1 [33, 50–52] and likely in meiosis as well. In addition, the interaction between CenpC and CenpA may stabilize them at centromeres [53] and may be strengthened by Aurora B-mediated phosphorylation.

Telomeres are best known for protecting chromosome ends from aberrant DNA damage responses. Intriguingly, telomeric and subtelomeric HC is broadly conserved [3, 54–56]; yet, HC is not directly involved in chromosome end protection under most conditions. This raises a key question: what selective advantage does telomeric HC confer? Our findings suggest that this conservation reflects a critical role in meiosis, where telomeric HC safeguards centromere identity. The telomere bouquet acts as a HC reservoir, enabling trans-acting reinforcement of pericentric HC and the localized activation of Aurora B kinase, which promotes centromere reassembly. In this way, telomeres safeguard not only chromosome ends, but also centromeres.

## Acknowledgements

We thank Eftychia Kyriacou for help with initial characterization of the *cnp1*-S5A alelle, Rahul Thadani for help with sequence alignment of CenpA and CenpC, and Rishi Nageshan for help with imaging analysis. We are grateful to Maria Diaz de la Loza for help with illustrations, and all our laboratory members for valuable discussions. We thank the Japanese National Bioresource Project for distributing strains harboring the *ark1.as*, *ark1*.T7, *hht1/3-T3A*, *hrk1*Δ, *pds5*Δ, *bir1-CD* and *hrk1*-CD alleles. This work was supported by NIGMS grant R01GM145820 and the University of Colorado School of Medicine.

## Author Contributions

HH performed the experiments presented in Figures 1,3,4,5,S1,S2,S3 and S4. ALM performed the experiments in Figures 2 and S1, and assisted with strain construction and experiments for Figures 1D, 5, S4D and S4E. YL assisted with imaging analysis for experiments presented in Figures 1A,3B,S1D,S2B and S4A. HH and JPC conceived the study, designed and interpreted the experiments, and wrote the manuscript. The authors declare no competing interests.

## Data Availability Statement

The raw imaging data that support the findings of this study are available upon request from the corresponding authors.

## Supplementary Figure Legends

**Figure S1.**
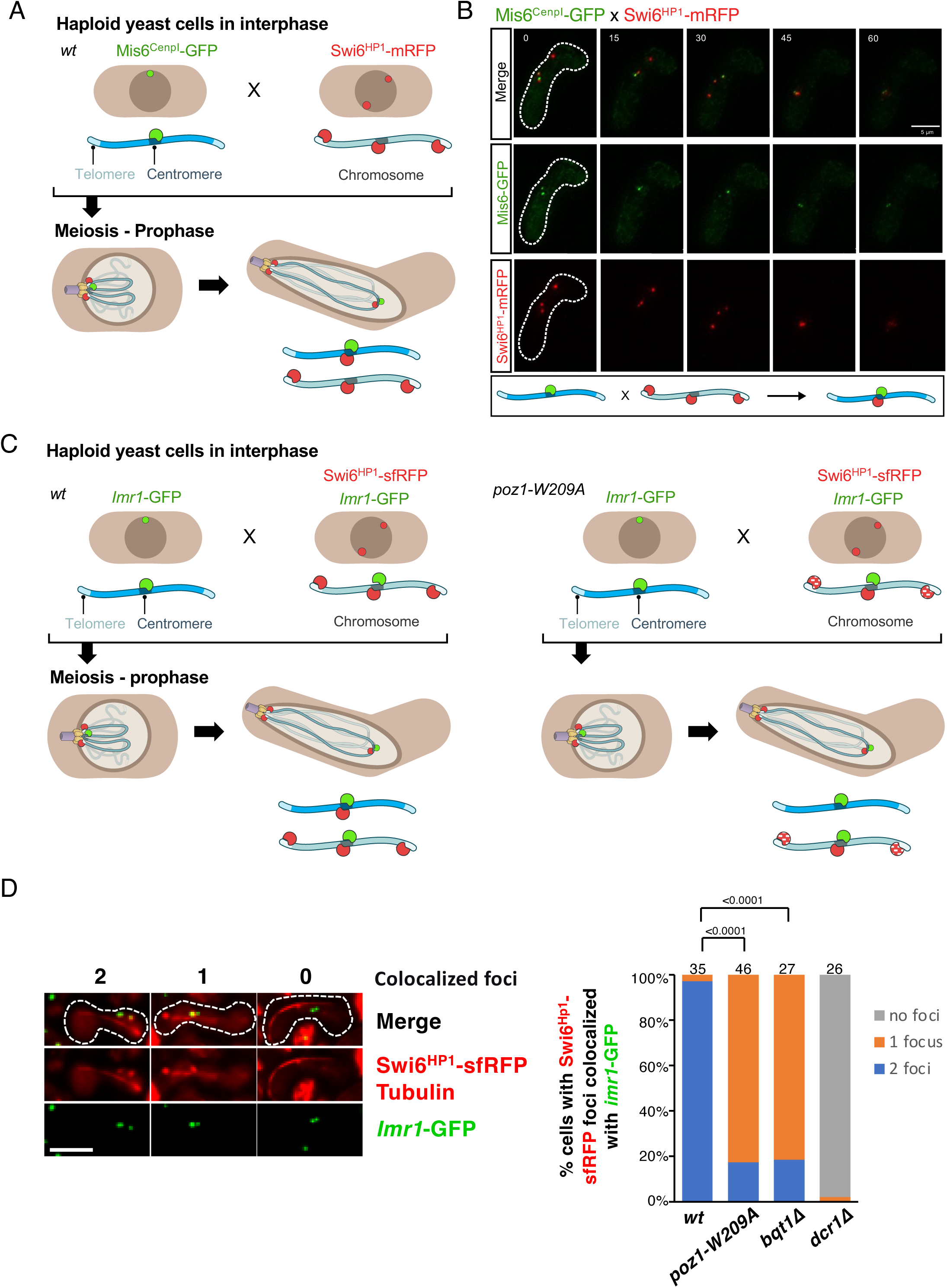
Telomeric heterochromatin promotes meiotic centromere assembly. **A.** Schematic of an additional experimental setup for visualizing pericentromeric HC assembly in meiosis. Haploid cells of opposite mating types, both harboring lacO/I-GFP arrays at the *Imr1* region of the pericentromere, are mated; only one mating partner expresses Swi6^HP1^-sfRFP. The diagram illustrates the *wt* scenario, in which Swi6^HP1^ is transferred from the Swi6^HP1^-sfRFP-expressing partner to promote pericentric HC assembly in the partner lacking sfRFP, resulting in colocalization of the both *Imr1*-GFP foci with Swi6-sfRFP in meiotic prophase. **B.** Examples and quantitation of Imr1-GFP foci colocalizing with Swi6^HP1^-sfRFP (categorized as colocalization in two, one, or no foci). Failure to transfer HC is observed in *poz1-W209A*, *bqt1Δ*, and *dcr1Δ* settings. The total number of meiocytes analyzed is indicated above each bar. P values, determined by two-tailed Fisher’s exact tests, are indicated above the brackets.

**Figure S2.**
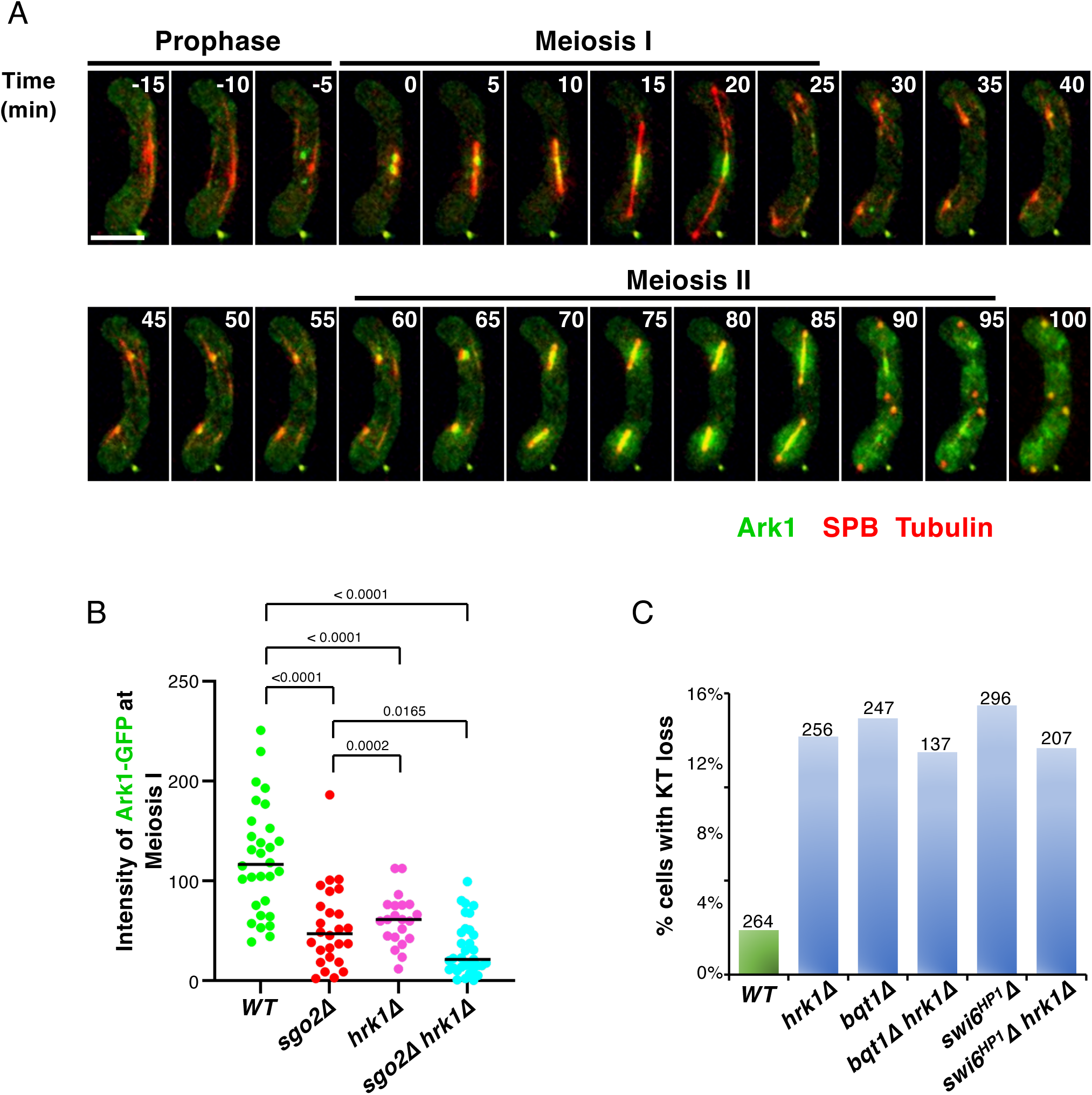
Ectopic CPC recruitment rescues meiotic centromere disassembly. **A.** Representative time-lapse images of CPC localization during meiosis. CPC (Ark1–GFP), SPBs (Sid4–mCherry) and microtubules (mCherry– Atb2^α-Tubulin^) are visualized. t = 0 is the first frame in meiosis I. Ark1-GFP foci are present before the onset of meiosis I (−5min). Scale bar = 5 μm. **B.** Quantitation of Ark1-GFP intensity along the spindles during metaphase I as exemplified in Figure 2B. Centromeric localization of Ark1-GFP is reduced in *sgo2Δ* and *hrk1Δ* single mutants and is further diminished in the *sgo2Δ hrk1Δ* double mutant. P values, determined by two-tailed Fisher’s exact tests, are indicated above the brackets. **C.** Epistasis analysis of mutants affecting meiotic centromere assembly. The *hrk1Δ* mutation does not enhance the KT loss phenotype observed in *bqt1Δ* or *swi6Δ* single mutant cells. The total number of meiocytes analyzed is indicated above each bar.

**Figure S3.**
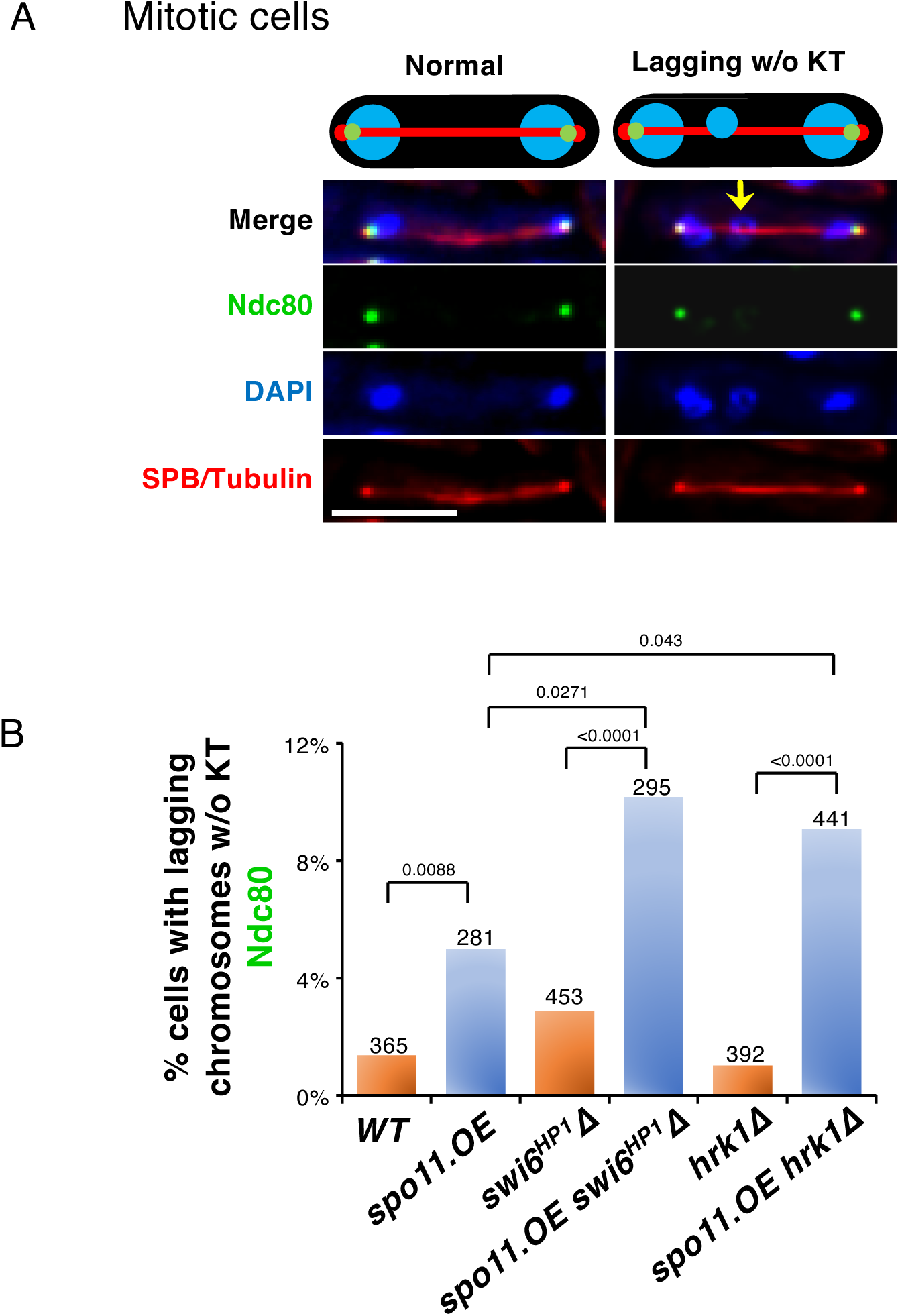
Compromised CPC exacerbates Spo11-mediated centromere dismantlement in proliferating cells. **A.** Representative images of KT loss in proliferating anaphase cells. Snapshots show fixed cells in late anaphase, with KT (Ndc80–2×GFP), chromatin (DAPI), SPBs (Sid4–mCherry) and spindles (mCherry–Atb2^α-Tubulin^) visualized. Arrows indicate lagging chromosomes lacking KTs. Scale bar = 5 μm. **B**. Quantitation of KT loss as in **A**. The KT loss rate, which is elevated by Spo11 expression, is further increased by loss of Swi6^HP1^ or Hrk1. The total number of anaphase cells analyzed is indicated above each bar. *P* values, determined by two-tailed Fisher’s exact tests, are indicated above the brackets.

**Figure S4.**
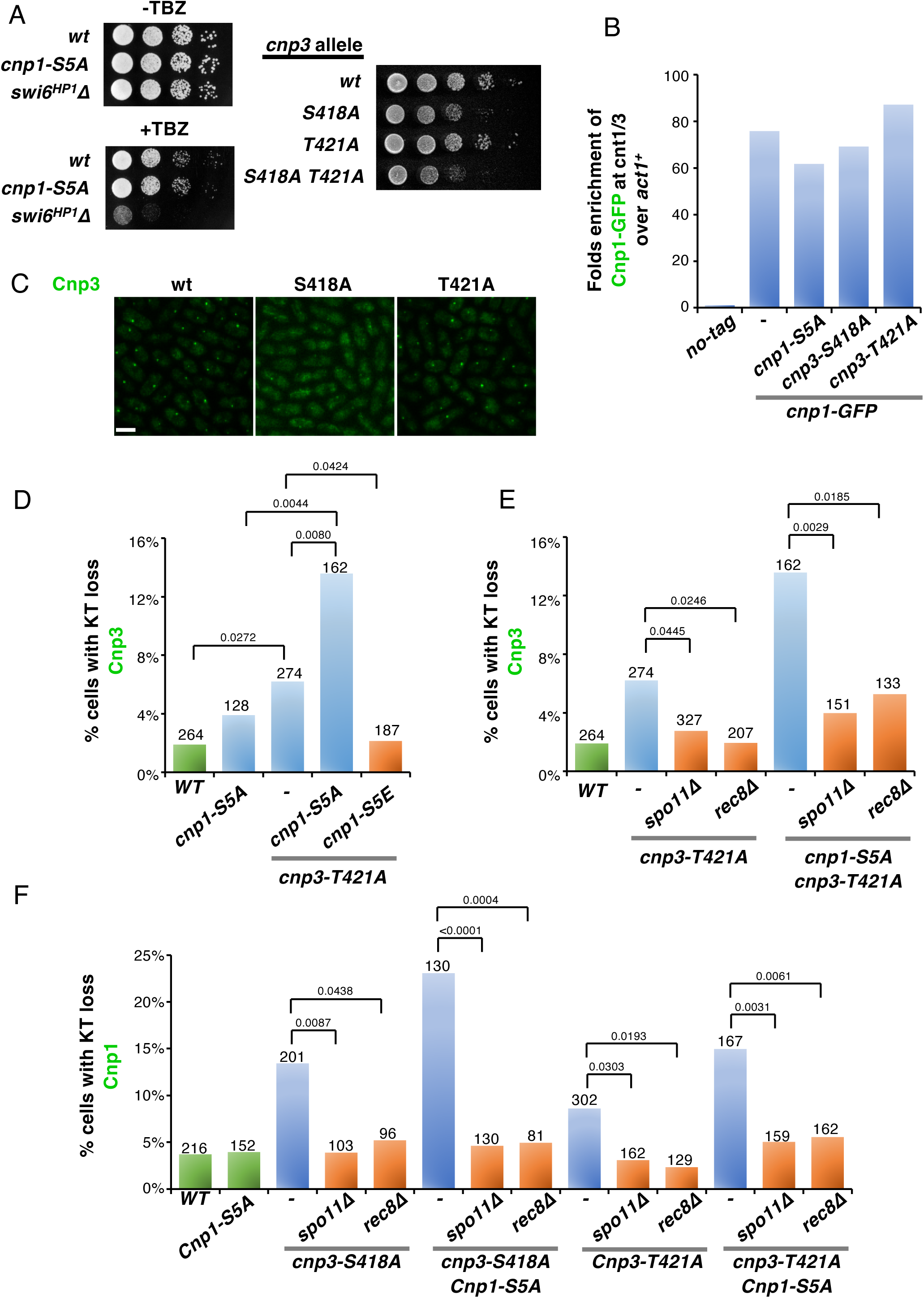
CenpA and CenpC phosphorylation by Aurora B promotes meiotic centromere assembly. **A.** Growth analysis of *cnp1* and *cnp3* phosphomutant alleles. The *cnp1-S5A* mutation does not confer growth defects in the absence or presence of TBZ. The *cnp3-S418A* mutation, but not *cnp3-T421A*, impairs cell growth; combining T421A with S418A results in a further growth impairment. **B.** ChIP analysis of Cnp1-GFP. *cnp1-S5A*, *cnp3-S418A*, and *cnp3-T421A* mutations do not significantly alter Cnp1 enrichment at the centromeric cores. **C.** Representative Images of Cnp3-GFP in asynchronous cultures of mitotically proliferating cells. The *cnp3-S418A* mutation reduces its centromeric localization. **D-F.** Quantitation of KT loss using Cnp3-GFP **(D–E)** or Cnp1-GFP **(F)** as the KT marker. The phosphomutation *cnp3-T421A* elevates meiotic KT loss, a phenotype that is further enhanced by the *cnp1-S5A* mutation **(D)**. The KT loss phenotypes in both the *cnp3-T421A* single mutant and *cnp3-T421A cnp1-S5A* double mutant are rescued by loss of Spo11 or Rec8 **(E)**. While *cnp1-S5A* alone does not increase KT loss, it enhances the KT loss rates of cells harboring *cnp3-S418A* and *cnp3-T421A* mutations. All of these KT loss phenotypes are rescued by *spo11Δ* or *rec8Δ* **(F)**. The total number of meiocytes analyzed is indicated above each bar. P values, determined by two-tailed Fisher’s exact tests, are indicated above the brackets.

## Materials and Methods

### Yeast strains and media

Supplementary Table 1 lists strain genotypes. The media used were as previously described ^1^. The rich medium YES, containing yeast extract, glucose, and five supplements (adenine, histidine, leucine, lysine, and uracil), was used for general cell proliferation. EMM5S (Edinburgh Minimal Medium with five supplements) was used for cell growth before imaging. EMM-N, which is identical to EMM5S but omits NH₄Cl to create a nitrogen-starvation medium, was used for imaging meiotic cells. SPA (Sporulation Agar), also a nitrogen-starvation medium, was used for mating and the induction of meiosis.

To generate point mutations encoding Cnp1-S5A, Cnp3-AA-GFP, and Cnp3-DD-GFP, one allele of the endogenous *cnp1+* or *cnp3+* gene in diploid cells was replaced with a PCR-generated mutant allele fused to a selectable marker (*hphMX* or *kanMX*). Haploid mutants were then isolated through meiosis and sporulation. Additional tagged versions of these mutants were later created using standard protocols with different selection markers ^2^. The Cnp1-S5E strain was constructed by replacing a *ura4+* marker at the endogenous *cnp1+* locus (*cnp1Δ::ura4+*) with the mutant allele, while a functional *cnp1-1* copy was maintained at the *lys1+* locus. This mutant was backcrossed to retain only the mutation at the endogenous locus. Other tagged versions were subsequently generated using the standard protocol. The strain containing *ark1.as* generated by the Yanagida laboratory was obtained from the Japanese National Bioresource Project (ID: FY20306), and sequencing confirmed that an *ark1-L166V* mutant allele was integrated into the endogenous locus. Other strains were obtained from the Japanese National Bioresource Project or as previously described ^1,3,4^. Strains with multiple mutated genotypes were generated through mating, sporulation, and selection for recombinants.

### Microscopy

Most microscopy images, except for those in Figures 1A, 3B, 4E, and Supplementary Figure S1D, S2B, were acquired on a DeltaVision microscope system (GE Healthcare Life Sciences). The system was equipped with an Olympus IX70 widefield inverted epifluorescence microscope, an Olympus UPlanSapo ×60 NA 1.42 oil immersion objective, a Photometrics CCD CoolSnap HQ camera, and an environmental chamber with a DeltaVision Spectris (GE Healthcare Life Sciences). Images were acquired with identical settings for light intensity, exposure time, number of focal planes, and step size within each experiment. All acquired image stacks were deconvolved and projected into two-dimensional maximum intensity projections using SoftWorx (GE Healthcare Life Sciences). Final image processing and analysis were performed using Fiji, an ImageJ-based open-source software.

Sample preparation and image acquisition protocols were as previously described (Hou et al., 2021), unless stated otherwise. Briefly, for live-cell imaging of meiosis, *h^90^* cells or a mixture of *h^+^* and *h^−^* cells were cultured and imaged at 30 °C throughout the experiments. Cells were first pre-grown overnight on EMM5S plates, spotted onto sporulation agar (SPA) plates, and incubated for 5–7 hours. They were then adhered to 35-mm glass-bottom culture dishes (MatTek) and immersed in 3 ml of nitrogen-deficient minimal medium (EMM-N) for microscopy. Time-lapse images were acquired in both mCherry and FITC (for GFP) channels over 22 focal planes at a 0.4 μm step size. To capture meiotic KT loss events, images were acquired at 15-minute intervals. A meiotic KT loss event was scored when one or more chromatin masses visualized with Pht1^H2AZ^–mCherry remained unsegregated and lacked detectable Cnp3-GFP foci. For time-lapse imaging of Ark1^AuroraB^–GFP localization during meiosis, images were acquired at 5-minute intervals. To image the co-localization of Swi6-sfRFP and Mis6^CenpI^-GFP during meiotic prophase, the characteristic morphologies of Swi6-sfRFP and Mis6^CenpI^-GFP foci were used to determine the cell cycle stage, and the extent of co-localization was quantified.

To capture KT loss events in mitosis, mid-log phase cells grown in YES media at 32°C were collected, fixed with ice-cold 70% ethanol for 15 minutes, centrifuged at 3,000 rpm for 30 seconds, resuspended in 1 ml of water for rehydration, and centrifuged again. The pellet was resuspended in 1 ml of PBS containing 5 μg/ml DAPI, incubated for 3 minutes, and washed three times with 1 ml of PBS. Images (2,048 × 2,048 pixels) were acquired across 25 focal planes with a 0.4 μm step size, and projected to 2D images after deconvolution. Cells were designated as being in late anaphase when the distance between the two SPBs (Sid4–mCherry) exceeded 6 μm. A mitotic KT loss event was scored when one or more DAPI-stained chromatin mass(es) remained unsegregated in late anaphase and lacked detectable Ndc80-2xGFP foci.

Imaging experiments for Figures 1A, 3B, 4E, and Supplementary Figure S1D, S2B were performed using a Nikon Ti2-LAPP system equipped with CFI SR HP Apo TIRF 60x and 100x oil immersion objectives, and the images were projected and analyzed using Fiji.

To examine telomere bouquet formation, meiotic cells were imaged on a Nikon Ti2-LAPP system using a sequential acquisition protocol for red fluorescence and FITC channels, followed by bright-field illumination. Z-stacks consisting of 7 optical sections were acquired at 0.6 µm intervals. Meiotic prophase cells were identified by the presence of mCherry-Atb2-labeled microtubule bundles emanating from a single Sid4-mCherry-labeled spindle pole body (SPB). Telomere bouquet formation was assessed using Taz1-GFP and scored when a single Taz1-GFP focus colocalized with the SPB and the Pht1^H2AZ^–mCherry-labeled chromatin mass was connected to the microtubule bundles via the SPB. In the absence of bouquet formation, multiple Taz1-GFP foci were observed, and the chromatin mass showed no connection to microtubules or the SPB.

To measure Ark1–GFP intensity at metaphase of meiosis I, *h^90^*cells were spotted on an SPA plate. After incubation at 28°C for 12 hours to allow mating and zygote formation, images were acquired on a Nikon Ti2-LAPP system using a sequential acquisition protocol for the red fluorescence and FITC channels, followed by bright-field illumination. Z-stacks of 15 optical sections at 0.4 µm intervals was collected. Cells in metaphase I were identified when the distance between the two SPBs (Sid4-mCherry foci) was less than 4 µm. A 7-pixel-wide line spanning the 4 µm distance through the two SPBs was drawn. The mean fluorescence intensity of Ark1-GFP along this line was measured, and the mean intensity of an identically sized line segment placed in a nearby cytoplasmic region of the same cell was subtracted as background.

To image the co-localization of Swi6-sfRFP and *Imr1*-GFP in meiotic prophase, cells were imaged on a Nikon Ti2-LAPP system using sequential acquisition for red fluorescence, FITC, and bright-field illumination. Z-stacks of 15 optical sections was acquired at 0.4 μm intervals. Meiotic prophase cells were identified by microtubule bundles (marked by mCherry-Atb2) emanating from a single SPB (marked by Sid4-mCherry).

To image Cnp3-GFP in proliferating mitotic cells, cells were grown to mid-log phase in liquid YES media, collected by centrifugation, resuspended in EMM5S media to an OD₅₉₅ of 0.2, grown further to OD_595_ 0.5–0.8 and harvested by centrifugation. Image acquisition was performed sequentially in the FITC and red fluorescence channels, followed by bright-field illumination on a Nikon Ti2-LAPP system. Z-stacks of 11 optical sections were acquired at 0.6 μm intervals.

### Chromatin immunoprecipitation (ChIP) and quantitative PCR (qPCR)

ChIP was performed using a standard protocol ^5^ with a few adjustments. Cells grown in YES liquid media at 30°C to mid-log phase (6 ×10^8^ cells in 50 ml) were crosslinked with 1% formaldehyde for 15 min at room temperature, harvested by centrifugation, and lysed by bead-beating in 400 μL lysis buffer (50 mM HEPES-KOH pH 7.5, 140 mM NaCl, 1 mM EDTA, 1% Triton X-100, 0.1% sodium deoxycholate) containing protease inhibitor cocktail (Beyotime) using a FastPrep-24 homogenizer (MP Biomedicals). Chromatin was sheared by sonication (Qsonica Q800R3) to 0.5–1.5 kb size range and immunoprecipitated with anti-GFP antibody (TransGen Biotech, HT801) and Protein A/G Resin (TransGen Biotech, DP501). After reversing crosslinking, immunoprecipitated DNA was purified with the GeneJET PCR Purification Kit (Thermo Scientific, K0702).

The qPCR protocol and primer sets were as previously described ^1^. Reactions were performed with ChamQ Blue Universal SYBR qPCR Master Mix (Vazyme, Q312) on a QuantStudio 3 system (Applied Biosystems). For each primer set, a standard curve was generated from serial dilutions of genomic DNA. Target sequence concentrations were calculated from these standard curves, with all reactions run on the same plate. Enrichment of target DNA was determined by normalization to whole cell extract and then to a reference fragment.

